# Functional Redundancy in Ocean Microbiomes Controls Trait Stability

**DOI:** 10.1101/2021.06.18.448980

**Authors:** Taylor M. Royalty, Andrew D. Steen

## Abstract

Theory predicts that functional redundancy in microbial communities increases trait stability, meaning that traits or functions are less likely to be lost from the community when species go extinct. However, few experiments have empirically tested this prediction, especially in the context of microbial communities and at the landscape scale. In part, the lack of metrics for functional redundancy in microbial ecosystems has prevented addressing this question. In a companion manuscript we proposed a quantitative metric for functional redundancy called Contribution Evenness (CE) that is optimized to reflect trait stability. Here, we use CE to predict the stability of marine microbial functions to species and transcription loss. Using transcriptomes deposited in the Ocean Microbial Reference Gene Catalog (OM-RGC.v2), a catalog of genes and transcripts sequenced by the TARA Ocean expedition, we quantified the functional redundancy for 4,314 KEGG Orthologs (KOs) across 124 marine sites. Functional redundancy was highly correlated with a latent variable consisting of four ocean physiochemical parameters: oxygen and chlorophyll a concentrations, depth, and salinity. Functional redundancy was higher at the poles than in non-polar regions. Simultaneously, regional β-diversity for individual functions was higher for functions with higher functional redundancy. These observations provide evidence that higher functional redundancy indicates increased stability of microbial ecosystem functions on spatiotemporal scales consistent with surface ocean mixing. We suggest that future changes in ocean physiochemistry could likely influence this stability for functions with lower functional redundancy.

**Importance:** Functional redundancy describes the state of multiple species performing the same function. Theory suggests functional redundancy stabilizes microbial community functions from disturbances leading to species loss or other changes to the microbiome. Previous work suggests that functional redundancy is common in ocean microbiomes which implies traits should be more stable among metacommunities. Some laboratory experiments demonstrate this idea, but it is difficult to test in the natural world. In a companion manuscript, we proposed a functional redundancy metric sensitive to trait stability. Here, we used this metric to show that functional redundancy varied substantially among ocean microbiomes and that regions with higher functional redundancy had higher regional trait stability. Last, we noted that variations in functional redundancy strongly correlated to ocean physiochemistry. Thus, changes in ocean physiochemistry via climate change may alter community traits to become more or less resistant to disturbance relative to contemporary conditions.

## Introduction

The ability to resolve complete genomes in microbial communities via contig binning and single-cell genome sequencing has revealed that metabolic functions, or traits, are often present in multiple taxa within a microbiome (1–4). The practical consequences of functional redundancy, in terms of community composition and function, are not well understood for microbial communities. Theoretical predictions, developed in the context of macroecosystems, suggest that functional redundancy can buffer a community against function loss due to species extinction (5–9). Foundational work on grassland ecosystems validated these theoretical predictions by showing that plant community biomass stability increased with higher functional redundancy (10). Mesocosm experiments provide empirical evidence that functional redundancy may also buffer microbial ecosystem functions from species loss. For instance, microbial density and biomass was less variable for assemblages with redundant populations occupying different trophic guilds in a food web (11), and reduction-oxidation conditions in sediment communities were more stable with higher bacterial diversity and niche overlap (12). Although existing theoretical and empirical evidence suggests functional redundancy may enhance ecosystem function stability, small-scale and mesocosm experiments with microbial communities may not reflect the most important processes on ecosystem scales (13). Establishing whether functional redundancy stabilizes traits is important for understanding how environmental fluctuations, most notably climate change, may influence microbially-mediated biogeochemical cycles.

In a companion paper, we developed a metric for functional redundancy called contribution evenness (CE) (14). CE uses genes or transcripts as traits (15, 16) and measures how evenly taxa contribute to the community-wide level of these traits (total gene abundance in the case of genes, and total transcript abundance in the case of transcripts). CE is optimized to reflect how stable microbially-catalyzed functions are to random species extinction or transcription loss in communities. Here, we use CE to address three fundamental questions about functional redundancy in marine microbial metacommunities:

1. Does functional redundancy of ocean microbiomes vary geographically
2. If so, does this variation relate to the physicochemical environment, and
3. What ecological consequences are apparent from observed patterns in functional redundancy?

To answer these questions, we used KEGG orthology (KO) annotations of genes reported from 180 sites sampled during the TARA Oceans expeditions, plus transcripts reported from a subset of 124 of those sites (17). KEGG is an attractive ontology because the functions of KOs are biochemically well-characterized and provide a reasonably large coverage of predicted genes in marine microbiomes. The *TARA Oceans* dataset is ideal for this work because it encompasses a diverse set of sites and marine ecosystems, which were sampled with generally standardized methods (18). By applying CE to the *TARA Oceans* data set, we identified geographic variations in functional redundancy, identified physicochemical factors relating to this variation, and established that microbial functions were less stable in regions with lower functional redundancy. These observations combined allowed us to identify some possible effects that climate change may have on the resilience of microbial community function to local disturbance.

## Methods

### TARA Oceans Genes and Environmental Data

We downloaded the entire Ocean Microbial Reference Gene Catalog v2 (OM-GRC.v2) on 1 Feb 2021 from EMBL-EBI (BioStudy: S-BSST297) and the environmental data from https://doi.org/10.5281/zenodo.3473199 (17). This data set includes annotations and abundance for metagenomes from 180 unique sites and transcriptomes from 124 sites. Gene and transcript profiles correspond to biological sequences collected from 100L of seawater on filters with size ranges 0.22-1.6 μm or 0.22-3.0 μm (18). Analysis was limited to sequences in OM-RGC.v2 annotated to KEGG orthologs (KO) (19). We interpreted length-normalized short-read mapping frequencies as abundances. These values are reflect the abundance of any sequence in a metagenome or transcriptome, given a fixed sequencing effort. Gene abundances for KO single copy marker genes (list below) annotated to the same genus were averaged to estimate the abundance of a given genus. Our selection of genus level analysis was to mitigate noise associated with analyzing functional redundancy of ASVs, which is theoretically possible. To allow for meaningful comparisons of functional redundancy at different sites, we rarified sequencing effort at each site. Relative abundances were then used as weights during random sampling, with replacement. Each site was sampled 3358334 and 472163 times for gene and transcript profiles, respectively. These values corresponded to the lowest sampling effort among all metagenomes and transcriptomes, respectively.

### KO Functional Redundancy and Diversity

Functional redundancy was calculated for each KO at each site for both metagenomes and metatranscriptomes. Functional redundancy was calculated with Contribution Evenness (CE) as detailed in the companion paper (14). Here, we treat KOs as proxies for metabolic traits (16, 20). In brief, the abundance of sequences sharing the same KO annotation in a sample was summed together, without regard to genus. Subsequently, the abundance of genes for individual genera was normalized by the summed abundance to obtain a relative abundance of each gene, for each genus. This vector corresponds to the relative contribution distribution (equation 3.2 in the companion paper). Species richness (*S*) used for calculating CE in equation 8 in companion paper was calculated with KO single copy marker genes. The presence of a genus was determined by summing the abundance of single copy marker genes: K06942, K01889, K01887, K01875, K01883, K01869, K01873, K01409, K03106, and K03110. If the summed abundance was greater than 0, then the genus was considered present in the metagenome/metatranscriptome.

### Global Modeling of Functional Redundancy

We sought to characterize spatial trends in the functional structure of microbial communities. To do this, we performed redundancy analysis using the rda function from the R package, vegan (21). In this analysis, the functional structure of microbial communities was treated as a response variable and physio-chemical variables were treated as predictor variables. For sites where a given KO was absent, the KO was assigned a value of zero. This situation occurred ~13.6% and 18.5% of the time for gene and transcript profiles, respectively. Low functional redundancy does not necessarily imply that a function is ecologically unimportant. This is particularly true for traits with low functional redundancy that strongly correlate to phylogeny. Such examples include *Thaumarchaeota* and diazotrophs which act as the primary nitrifiers and nitrogen fixers in the ocean, respectively (22). On the contrary, small changes in functional redundancy for traits with low functional redundancy might reflect key ecological processes, and thus, comparisons of community functional structure should account for the differences in functional redundancy magnitude. Thus, changes in KO functional redundancy reflected changes with respect to the KO’s observed variance while not reflecting the KO’s order of magnitude, which spanned five orders of magnitude. To prevent functions with large CE from dominating the redundancy analysis, we normalized CE for each KO at each site by the respective KO’s variance from across all sites. For the redundancy analysis, we used salinity (PSU), nitrate (mmol m^−3^), phosphate (mmol m^−3^), oxygen (mmol m^−3^), chlorophyll A (mg m^−3^), depth (m), and silicate (mmol m^−3^). We chose these variables as they were *in situ* measurements (18) and could scale with models that predict global ocean chemistry. We imputed missing data, as removing incomplete cases can bias datasets (23). Random forest imputation was performed using the missForest function, from the R package, missForest (24). Next, each environmental variable was converted into a normal distribution using a boxcox transformation via the boxcox function in the R package, MASS (25). Variables were then centered to have a mean of zero. Last, the significance of each variable as well as the first two canonical axes was verified using the anova.cca function in the R Package, vegan (21).

The first canonical axis derived from the redundancy analysis was used to predict mean transcript functional redundancy from the individual metatranscriptomes (sites) using OLS regression. This model was compared to an OLS regression using salinity, nitrate, phosphate, oxygen, chlorophyll A, depth, and silicate as predictors of mean transcript functional redundancy. Similar to before, missing data were imputed using missForest (24), variables were transformed with a boxcox transformation (25), and variables centered so the mean distribution was zero prior to regression. The best predictor subset was determined using the regsubsets function from the R package, leaps (26). The criteria defining the best model was minimizing AIC.

The monthly-averaged data products spanning from Jan 2013 to Dec 2018 for GLOBAL_REANALYSIS_BIO_001_029 and GLOBAL_REANALYSIS_PHY_001_031 were downloaded from https://marine.copernicus.eu/ on 7 May 2021. Data products had a grid resolution of 0.25°x0.25°. The OLS regression using the first canonical axis as a predictor substantially outperformed the best subset OLS when predicting mean transcript functional redundancy. We therefore converted the predicted data product chemistry for each grid cell into a score for the first canonical axis. Then, the mean transcript functional redundancy was predicted for each grid cell, for each month, utilizing the coefficient derived from the canonical axis OLS regression. The median, 5^th^ percentile, and 95^th^ percentile of mean functional redundancy was taken across the six-year window. Variance was measured as the difference between the 95^th^ and 5^th^ percentile divided by the median.

### Metatranscriptome β-diversity and Trait Stability

The *β*-diversity (27) was calculated for individual functions (KO) across different regions. Regions were defined as a metatranscriptome and the nine closest metatransciptomes in the epipelagic (depth < 200m). Site distances were measured using the published site coordinates (17) and the *distm* (default parameters) function from the R package, geosphere (28). Redundant combinations of sites were removed prior to reporting results. The average functional redundancy for individual functions was taken for each region. The *β*-diversity was calculated as:

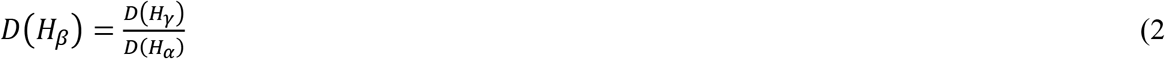

Here, we used a diversity order q=0, or the richness to calculate diversity. As such, *γ*-diversity was either 0 (regionally absent) or 1 (regionally present), and *α*-diversity was the proportion of local communities with the trait. Since *β*-diversity is not defined for traits completely absent from a region, the *β*-diversity calculation simplified to:

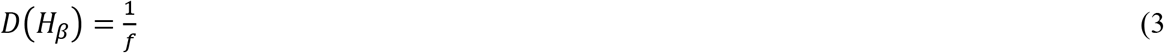

which is simply the inverse proportion (*f*) of local communities in a region with a given function.

## Results and Discussion

We used CE (14) to measure functional redundancy for KEGG ortholog (KO)-annotated genes from 180 and 124 *TARA Oceans* metagenomes and metatranscriptomes, respectively (17). CE uses genes (CE with respect to genes) or transcripts (CE respect to transcripts) as traits (15, 16) and measures how evenly species in a community contribute to these traits as a whole. After calculating CE for metagenomes (Fig. 1A) and metatranscriptomes (Fig. 1B), we found that KOs spanned over three orders of magnitude, demonstrating the high degree of variability in how functionally redundant traits are in the ocean. We further isolated KOs based on KEGG energy metabolism classification to illustrate how functional redundancy varied across the ocean for sulfur, photosynthesis, oxidative phosphorylation, nitrogen, methane, and carbon fixation. The distribution of transcript abundance within ocean microbiomes correlated well with the distribution of gene abundance at the same site (Fig. 1C; R^2^=0.45, p<0.001). This relationship was not surprising considering previous reports that metagenome gene abundances strongly influenced transcript abundances across *TARA Oceans* microbiomes (17). Although the nature of these observations is similar, the ecological implication is different. Specifically, the observation of Salazar et al. shows correlation in absolute gene abundance and transcription abundance, whereas our observation indicates that when genomic traits are more evenly distributed across genus within a community, transcription is also more evenly distributed across genus.

**Fig. 1:**
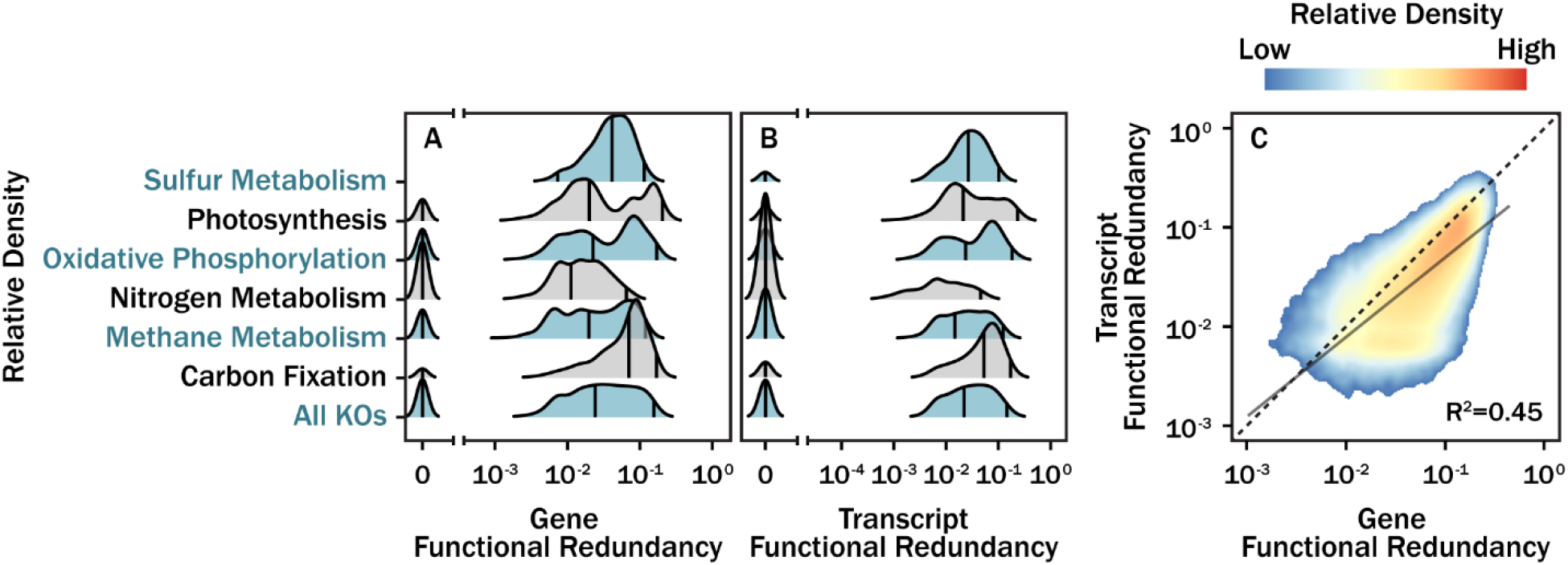
Variation in functional redundancy for KOs across 180 *TARA Oceans* metagenomes (**A**), 129 *TARA Oceans* transcriptomes (**B**), and a pairwise comparison of functional redundancy (*n*≈415,000) for KOs annotated the same at the same site (**C**). Density curves (**A,B**) were generated with a kernel density estimate (gaussian kernel with a bandwidth 0.0803). The vertical black lines correspond to 5^th^, 50^th^, and 95^th^ quantiles. The metagenomes and metatranscriptomes functional redundancy regression (**C**) followed a log10-log10 model (solid black line). Prior to regression, all values were offset by 10^−4^ and high leverage data was removed based on high Cook’s distance 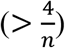. The color gradient and black dashed line correspond to data density and a one-to-one relationship, respectively.

To better understand the relationship between environmental characteristics and functional redundancy, we used redundancy analysis (RDA) to model the functional redundancy of each KO at each site as a function of the seven environmental parameters: salinity, depth, and concentrations of nitrate, phosphate, and silicate ions, oxygen, and chlorophyll a (Fig. 1). The first canonical axis of this model explained 19.8% of the total functional redundancy variance in 4,314 transcripts annotated as KOs. An ANOVA demonstrated that the first canonical axis explained significantly more variability in KO functional redundancy than a null model (*p*<0.01) (29). Although the total explanatory power for individual KOs was low, we found that the average functional redundancy of all KOs at each TARA Oceans site was accurately predicted by the first canonical axis alone (93.2% of total variance explained; ordinary least squares regression.

Factor loadings of this first canonical axis were dominated positively by oxygen and chlorophyll a and negatively by sample depth and salinity (Fig. 2B). Thus, this canonical axis appears to be a proxy for phototroph biomass, or the extent to which an environment is dominated by copiotrophs versus oligotrophs. The strong predictive power of chlorophyll a was not surprising given that carbon export mediated by primary productivity is known to be a good predictor for community composition and genomic composition in the ocean (30). Higher functional redundancy was positively correlated with this proxy for phototroph biomass. We compared the performance of our factor-based model to a best subset (based on an AIC criteria) OLS regression using the original seven physiochemical variables (NO_3_^−^, PO_4_^3-^, salinity, depth, O_2_, Si, and ChlA) that generated our factor (Fig. 2C). In contrast to the factor model, the best subset model (O_2_, Si, and depth) explained only 32.4% of the variance in mean functional redundancy.

**Fig. 2:**
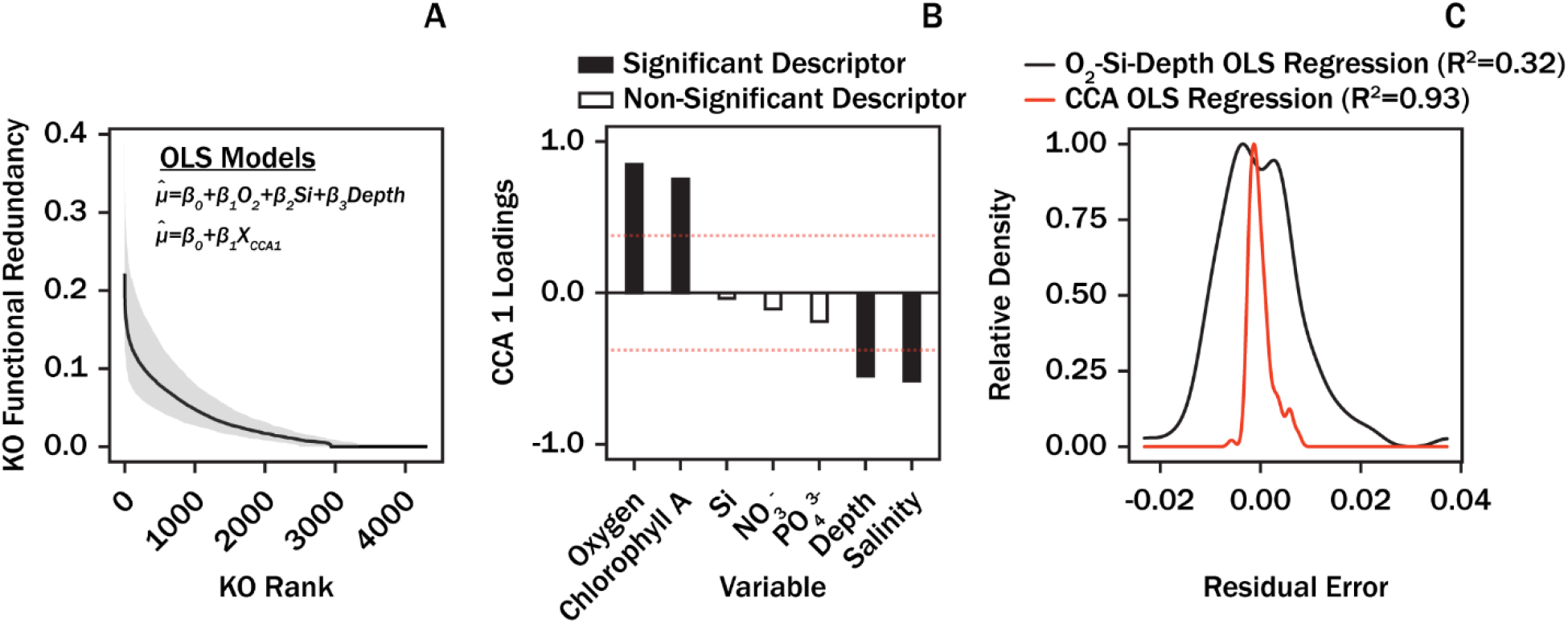
Metatranscriptome rank-functional redundancy curves across all *TARA Oceans* sites (n=129) analyzed in this study (**A**). The black line corresponds to the median while the shaded area corresponds to the range spanning between the 5^th^ and 95^th^ percentiles. Proposed models were evaluated for accuracy in predicting mean (*μ*) and standard deviation (*σ*) of individual rank-functional redundancy curves. The O_2_-Si-Depth model was selected as a best subset (minimum AIC) among the seven predictor variables. Canonical axis 1 loadings (scaling = 0) derived from the redundancy analysis (**B**). Dashed red lines in panel (**B)**correspond to the “equilibrium line of descriptors” 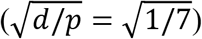, a threshold for defining significant variable contribution to factor loadings (36). A comparison in residual error of mean functional redundancy predicted by the O_2_-Si-Depth OLS regression model and the canonical axis OLS regression model (**C**).

The high explanatory power by the environmental data justified extrapolating our factor model onto a global scale. Using global predictions of ocean nutrient concentrations, we calculated factor scores at a 0.25°x0.25° resolution and predicted functional redundancy across Earth’s oceans using the derived coefficient from our OLS regression of mean functional redundancy versus the first canonical axis of the RDA. Our model predicts that functional redundancy is highest in polar regions and near river outflows, and lowest in subtropical gyres (Fig. 3A). Variance was highest in polar regions, coastal regions, and river outflows (Fig. 3B). The transition from high to low functional redundancy between polar and non-polar latitudes was consistent with previously reported ecological boundaries for ocean microbiomes, where the transition from non-polar to polar latitudes corresponded to compositional changes in metatranscriptomes and metagenomes (17).

**Fig. 3:**
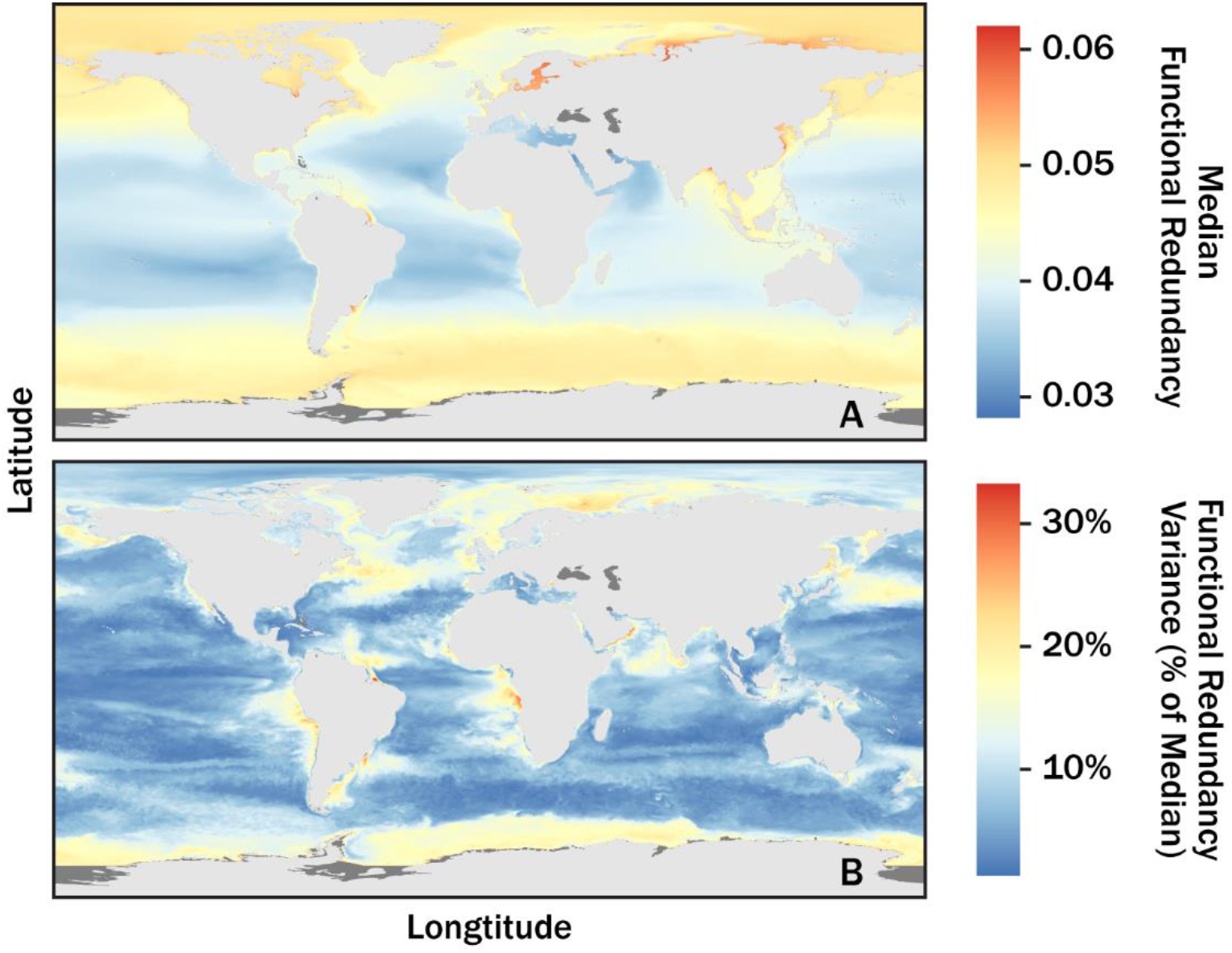
Mean functional redundancy of all KOs at a 5m depth across Earth’s oceans (0.25° × 0.25° resolution). Panels (**A**) and (**B**) correspond to the median and variance of predictions spanning Jan 2013 to Dec 2018, respectively. Dark and light grey correspond to regions absent of predictor data and land, respectively.

### Functional Redundancy as an Indicator of Trait Stability

In the companion paper, we presented a numerical simulation demonstrating a relationship between CE and trait stability to random extinctions. Here, we sought to establish whether there is empirical evidence that functional redundancy increases trait stability in ocean microbiomes, as previously hypothesized (31). Specifically, we hypothesized that if higher levels of functional redundancy do indeed increase trait stability, then the sets of functions present in nearby communities should be more similar when functional redundancy is higher. To test this hypothesis, we compared functional redundancy at each epipelagic site to the β-diversity (27) of each function (coded as present or absent) in a local region consisting of the site in question plus the nine nearest sites. In this context, lower β-diversity indicates that a larger portion of local communities shared a given function (equation 3). Since microbial communities exhibit spatial autocorrelation (32), these ad-hoc local regions serve as a reasonable alternative to the ideal data set, which would consist of subsequent metagenomes taken from a parcel of water over time.

We observed a negative monotonic relationship between functional redundancy and beta diversity for every KO for every unique region (Fig. 4). This qualitative evidence is consistent with the idea that functional redundancy maintains trait stability on a regional level. Notably, CE>0.01 corresponded to functions becoming almost entirely stable within their respective regions (Fig. 4). This threshold suggests a large portion of functions should be stable across Earth’s ocean microbiome (Fig. 3). Nonetheless, select functions do appear vulnerable to regional instability. Notably, nitrogen metabolism functions tend to have CE<0.01 (Fig. 1B). Climate models and historical observations indicate that major ocean currents are slowing (33), oxygen minimum zones are expanding (34), and there will be future changes in regional primary productivity (35). With respect to oxygen and primary productivity, these variables are important in our factor model and suggest that future changes in these parameters might disproportionately influence the stability for lower functional redundancy functions such as nitrogen energy metabolism.

**Fig. 4.**
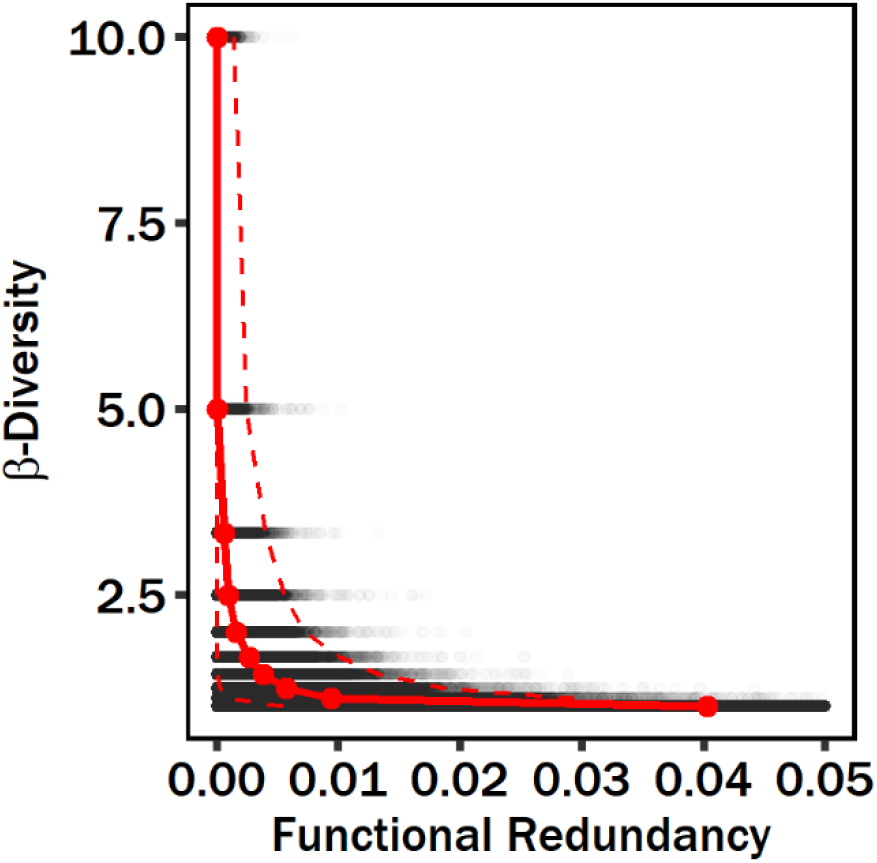
The *β*-diversity for individual KOs as a function of the average functional redundancy within local regions. The local region of each site was defined as the site plus the 9 geographically nearest sites. For all regions, the median distance from the region center to sites within the region is 1740 km. The red points and solid red line correspond to the median functional redundancy across all KOs for a given *β*-diversity. The upper and lower dashed red lines correspond to the 97.5 and 2.5 quantiles, respectively.

## Acknowledgements

Funding for this project was provided by the Department of Energy, Office of Science, Office of Biological and Environmental Research (DE-SC0020369) to A.D.S.

## Data and Materials Availability

The Ocean Microbial Reference Gene Catalog v2 (OM-GRC.v2) is accessible from EMBL-EBI (BioStudy: S-BSST297). *TARA Oceans* site metadata was downloaded from https://doi.org/10.5281/zenodo.3473199. The hindcast data products, GLOBAL_REANALYSIS_BIO_001_029 and GLOBAL_REANALYSIS_PHY_001_031, are accessible from https://marine.copernicus.eu/. All code and processed data is available at https://github.com/taylorroyalty/tara_ocean_fr.

## References and Notes

1. Louca S, Polz MF, Mazel F, Albright MBN, Huber JA, O’Connor MI, Ackermann M, Hahn AS, Srivastava DS, Crowe SA, Doebeli M, Parfrey LW. 2018. Function and functional redundancy in microbial systems. Nat Ecol Evol 2:936–943.

2. Royalty TM, Steen AD. 2019. Quantitatively Partitioning Microbial Genomic Traits among Taxonomic Ranks across the Microbial Tree of Life. mSphere 4.

3. Martiny JBH, Jones SE, Lennon JT, Martiny AC. 2015. Microbiomes in light of traits: A phylogenetic perspective. Science (80-) 350:aac9323.

4. Delmont TO, Quince C, Shaiber A, Esen ÖC, Lee STM, Rappé MS, McLellan SL, Lücker S, Eren AM. 2018. Nitrogen-fixing populations of Planctomycetes and Proteobacteria are abundant in surface ocean metagenomes. Nat Microbiol 3:804–813.

5. Naeem S. 1998. Species Redundancy and Ecosystem Reliability. Conserv Biol 12:39–45.

6. Rosenfeld JS. 2002. Functional redundancy in ecology and conservation. Oikos 98:156–162.

7. Fonseca CR, Ganade G. 2001. Species functional redundancy, random extinctions and the stability of ecosystems. J Ecol 118–125.

8. Walker BH. 1992. Biodiversity and Ecological Redundancy. Conserv Biol 6:18–23.

9. Cottingham KL, Brown BL, Lennon JT. 2001. Biodiversity may regulate the temporal variability of ecological systems. Ecol Lett 4:72–85.

10. Tilman D, Reich PB, Knops JMH. 2006. Biodiversity and ecosystem stability in a decade-long grassland experiment. Nature 441:629–632.

11. Naeem S, Li S. 1997. Biodiversity enhances ecosystem reliability 390:507–509.

12. Hunting ER, Vijver MG, van der Geest HG, Mulder C, Kraak MHS, Breure AM, Admiraal W. 2015. Resource niche overlap promotes stability of bacterial community metabolism in experimental microcosms. Front Microbiol 6:1–7.

13. ZoBell CE, Anderson DQ. 1936. Observations on the multiplication of bacteria in different volumes of stored sea water and the influence of oxygen tension and solid surfaces. Biol Bull 71:324–342.

14. Royalty TM, Steen AD. 2021. Contribution Evenness: A functional redundancy metric for microbially-mediated biogeochemical rates and processes. bioRxiv 6.

15. Violle C, Navas M-L, Vile D, Kazakou E, Fortunel C, Hummel I, Garnier E. 2007. Let the concept of trait be functional! Oikos 116:882–892.

16. Green JL, Bohannan BJM, Whitaker RJ. 2008. Microbial biogeography: from taxonomy to traits. Science (80-) 320:1039–1043.

17. Salazar G, Paoli L, Alberti A, Huerta-Cepas J, Ruscheweyh HJ, Cuenca M, Field CM, Coelho LP, Cruaud C, Engelen S, Gregory AC, Labadie K, Marec C, Pelletier E, Royo-Llonch M, Roux S, Sánchez P, Uehara H, Zayed AA, Zeller G, Carmichael M, Dimier C, Ferland J, Kandels S, Picheral M, Pisarev S, Poulain J, Acinas SG, Babin M, Bork P, Boss E, Bowler C, Cochrane G, de Vargas C, Follows M, Gorsky G, Grimsley N, Guidi L, Hingamp P, Iudicone D, Jaillon O, Kandels-Lewis S, Karp-Boss L, Karsenti E, Not F, Ogata H, Pesant S, Poulton N, Raes J, Sardet C, Speich S, Stemmann L, Sullivan MB, Sunagawa S, Wincker P. 2019. Gene Expression Changes and Community Turnover Differentially Shape the Global Ocean Metatranscriptome. Cell 179:1068–1083.e21.

18. Pesant S, Not F, Picheral M, Kandels-Lewis S, Le Bescot N, Gorsky G, Iudicone D, Karsenti E, Speich S, Trouble R, Dimier C, Searson S. 2015. Open science resources for the discovery and analysis of Tara Oceans data. Sci Data 2:1–16.

19. Kanehisa M, Furumichi M, Tanabe M, Sato Y, Morishima K. 2017. KEGG: New perspectives on genomes, pathways, diseases and drugs. Nucleic Acids Res 45:D353–D361.

20. Krause S, Le Roux X, Niklaus PA, Van Bodegom PM, Lennon JT, Bertilsson S, Grossart H-P, Philippot L, Bodelier PLE. 2014. Trait-based approaches for understanding microbial biodiversity and ecosystem functioning. Front Microbiol 0:251.

21. Philip D. 2003. Computer program review VEGAN, a package of R functions for community ecology. J Veg Sci 14:927–930.

22. Pajares S, Ramos R. 2019. Processes and Microorganisms Involved in the Marine Nitrogen Cycle: Knowledge and Gaps. Front Mar Sci 0:739.

23. McElreath R. 2020. Statistical Rethinking: A Bayesian Course with Examples in R and Stan, 2nd ed. CRC Press.

24. Stekhoven DJ, Bühlmann P. 2012. Missforest-Non-parametric missing value imputation for mixed-type data. Bioinformatics 28:112–118.

25. Venables WN, Ripley BD. 2002. Modern Applied Statistics with S. 7.3.49. Springer, New York.

26. Lumley T. 2004. The leaps Package. 2.7.

27. Jost L. 2007. Partitioning diversity into independant alpha and beta components. Ecology 88:2427–2439.

28. Hijmans R, Williams E, Vennes C. 2015. Geosphere: spherical trigonometry. R package. 1.5.

29. Legendre P, Oksanen J, ter Braak CJF. 2011. Testing the significance of canonical axes in redundancy analysis. Methods Ecol Evol 2:269–277.

30. Guidi L, Chaffron S, Bittner L, Eveillard D, Larhlimi A, Roux S, Darzi Y, Audic S, Berline L, Brum JR, Coelho LP, Espinoza JCI, Malviya S, Sunagawa S, Dimier C, Kandels-Lewis S, Picheral M, Poulain J, Searson S, Stemmann L, Not F, Hingamp P, Speich S, Follows M, Karp-Boss L, Boss E, Ogata H, Pesant S, Weissenbach J, Wincker P, Acinas SG, Bork P, De Vargas C, Iudicone D, Sullivan MB, Raes J, Karsenti E, Bowler C, Gorsky G. 2016. Plankton networks driving carbon export in the oligotrophic ocean. Nature 532:465–470.

31. Lennon JT, Jones SE. 2011. Microbial seed banks: The ecological and evolutionary implications of dormancy. Nat Rev Microbiol 9:119–130.

32. Ghiglione J-F, Galand PE, Pommier T, Pedrós-Alió C, Maas EW, Bakker K, Bertilson S, Kirchmanj DL, Lovejoy C, Yager PL, Murray AE. 2012. Pole-to-pole biogeography of surface and deep marine bacterial communities. Proc Natl Acad Sci U S A 109:17633–8.

33. Caesar L, McCarthy GD, Thornalley DJR, Cahill N, Rahmstorf S. 2021. Current Atlantic Meridional Overturning Circulation weakest in last millennium. Nat Geosci 14:118–120.

34. Ito T, Minobe S, Long MC, Deutsch C. 2017. Upper ocean O2 trends: 1958–2015. Geophys Res Lett 44:4214–4223.

35. Fu W, Randerson JT, Keith Moore J. 2016. Climate change impacts on net primary production (NPP) and export production (EP) regulated by increasing stratification and phytoplankton community structure in the CMIP5 models. Biogeosciences 13:5151–5170.

36. Legendre P, Legendre L. 2012. Numerical Ecology, 3rd ed. Elsevier.

